# OP-GLX: A MATLAB Toolbox for Online Processing and Plotting of Neuropixels Data Acquired with SpikeGLX

**DOI:** 10.64898/2026.03.04.709636

**Authors:** Jacob C. Slack, Gus Rutledge, Madison D. Doss, Amol P. Yadav

**Affiliations:** Lampe Joint Department of Biomedical Engineering, University of North Carolina at Chapel Hill and North Carolina State University, Chapel Hill, NC 27599, USA

## Abstract

Online processing and visualization of large-scale neural data is critical for neuroscientific discovery and advancements in neural engineering. However, with the development of technologies like Neuropixels (NP) probes, which enable simultaneous streaming from hundreds of recording electrodes, handling such data in real-time has become an ongoing challenge. Moreover, keeping pace with recording hardware has required most existing software, such as SpikeGLX for NP probes, to prioritize acquisition stability, leaving data processing and visualization to primarily be performed offline. Thus, we created OP-GLX, a MATLAB-based toolbox designed to operate in tandem with SpikeGLX to enhance the fetching, processing, and visualization of incoming neural data. The OP-GLX toolbox features several processing capabilities, including spike detection, computing time-binned firing rates, plotting spike waveforms, and conducting principal component analysis (PCA). The processed neural data is displayed on a native graphical user interface (GUI) for intuitive and customizable interaction with the experiment. The performance testing of OP-GLX showed that it supports real-time operation, confirmed by the absence of SpikeGLX stream buffer fetch errors across multiple acquisition settings. By complementing current neural data acquisition methods and providing stable online functionality, we envision that OP-GLX will enable researchers to visualize and interpret their data more effectively during ongoing neuroscience experiments.

## Introduction

Recent hardware advancements in semiconductor technology have promoted the creation of high-density recording devices with high spatial and temporal resolution. Such advancements include the Neuropixels (NP) probes, which are capable of simultaneously recording large populations of neurons across a linear, one-centimeter path [1]. These probes have become a standard tool for neuroscientists and are increasingly attractive for neural engineering and brain-computer interface (BCI) applications requiring large-scale recordings [2], [3]. The high spatial and temporal resolution of these probes has facilitated numerous scientific discoveries from both chronic [4], [5] and acute [6] studies. However, the data acquired from such probes is inherently high-dimensional and frequently generates data on the order of Gigabytes/minute [7], making visualization of incoming neural signals a challenge.

The ability to visualize high-dimensional neural datasets during data acquisition is crucial for neurological discovery, as it can facilitate a deeper understanding of the collected data.

Visualization tools also provide researchers with feedback on incoming data, enabling insights into variables such as probe placement, noise levels and stimulus features during data collection. Moreover, high-dimensional neural data streams are spatially [8] and temporally [9] complex, as features of interest can occur on sub-second timescales and across a multitude of neurological structures.

The current standard for NP data acquisition (DAQ) necessitates the use of either SpikeGLX [10] or Open Ephys [11] hardware/software platforms. While Open Ephys offers increased modularity and visualization of DAQ workflows, SpikeGLX was explicitly designed for stable, low-latency acquisition and recordings of high channel-count NP probes. Although widely adopted and robust, online data processing and visualization capabilities of SpikeGLX are largely limited to raw or filtered voltage traces and manual thresholding of single channels for spike detection with more complex analysis methods, such as spike-sorting, performed offline (Fig. 1A). Additionally, spike detection across all channels involves a single threshold value and is viewed on a heatmap of the probe, limiting the ability to visualize the relationships of activity among multiple spatial recording regions. Presently, few systems exist that provide advanced real-time processing methods or are highly specific to perform one task, such as spike-sorting [12]. Because of the increasingly widespread adoption of high-density NP probes for neuroscientific research and their relevance in BCI applications, there is an urgent need for ubiquitous systems that provide online data processing and visualization of large-scale neural recording streams.

**Figure 1:**
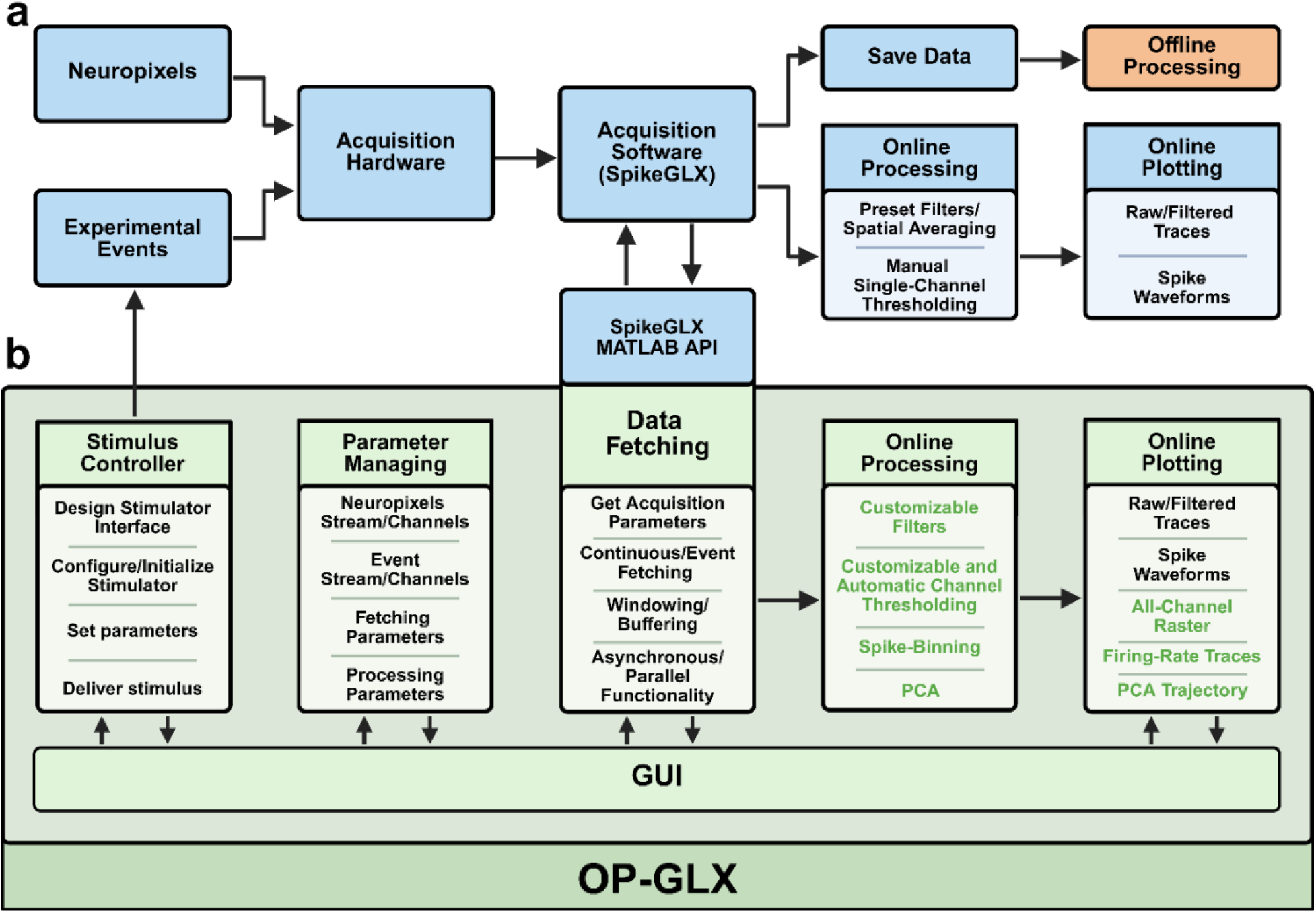
Overview of OP-GLX toolbox integration and features. **a**) Standard workflow of Neuropixels data acquisition via SpikeGLX. Priority is given to maintaining stable, low-latency, high-throughput acquisition where online analysis is limited and primarily performed offline. **b**) OP-GLX toolbox runs alongside SpikeGLX providing advanced functionality without impeding acquisition. OP-GLX utilizes asynchronous and parallel functionality to fetch data from SpikeGLX enabling complex processing methods (green text) to be performed online.

Here, we present OP-GLX: a MATLAB toolbox for online processing and plotting (OP) of NP data acquired by SpikeGLX. The primary purpose of OP-GLX is to neatly package incoming chunks of data from the 30kHz action potential (AP)-band streams utilizing the SpikeGLX MATLAB software development kit (SDK) [13] to allow for real-time data processing and visualization (Fig. 1B, Supplementary Video 1). Designed as a MATLAB toolbox, OP-GLX can be easily installed and incorporated within SpikeGLX acquisition workflows, where fetched data is processed and subsequently plotted in a separate MATLAB graphical user interface (GUI).

Additionally, OP-GLX design philosophy prioritizes both ease of implementation for non-technical users while offering the ability to integrate custom scripts and functions for advanced users. Source code, toolbox installation and instructions can be found on the GitHub repository: https://github.com/yadavlabs/OP-GLX.

### Toolbox Organization and Architecture

Development and testing of OP-GLX was performed in MATLAB versions R2024b and R2025b on a Windows 11 laptop with 16GB of RAM and a Windows 11 desktop with 128GB of RAM. SpikeGLX versions v20241215-phase30 (now removed) and v20250325-phase30 were used alongside the precompiled application programming interface (API) from the SpikeGLX-MATLAB-SDK (commit 26ae92d). SpikeGLX and OP-GLX instances were run on the same device with the SpikeGLX remote command server set to the default loopback address (127.0.0.1) and port (4142).

OP-GLX is a MATLAB toolbox built around the *SpikeGL* API class provided by the SpikeGLX-MATLAB-SDK. The toolbox structures and extends API calls enabling real-time fetching, processing, and visualization of NP acquisition streams during ongoing experiments. To avoid ambiguity, functions and methods originating from the SpikeGLX-MATLAB-SDK are referred to throughout the paper as the “API” or “API calls”; OP-GLX-originating functionality will be referred to explicitly by class, method, or function name. The precompiled *SpikeGL* API is packaged with the toolbox and can be found in /toolbox/external/ directory of the repository.

The OP-GLX toolbox is organized using MATLAB namespaces (e.g., +packageName) to group functions and classes by functionality while preventing name collisions with user code or third-party packages (Table 1). Core components include data fetching and buffering, parallel processing, spike detection and feature extraction, and GUI-based control and visualization.

**Table 1:**
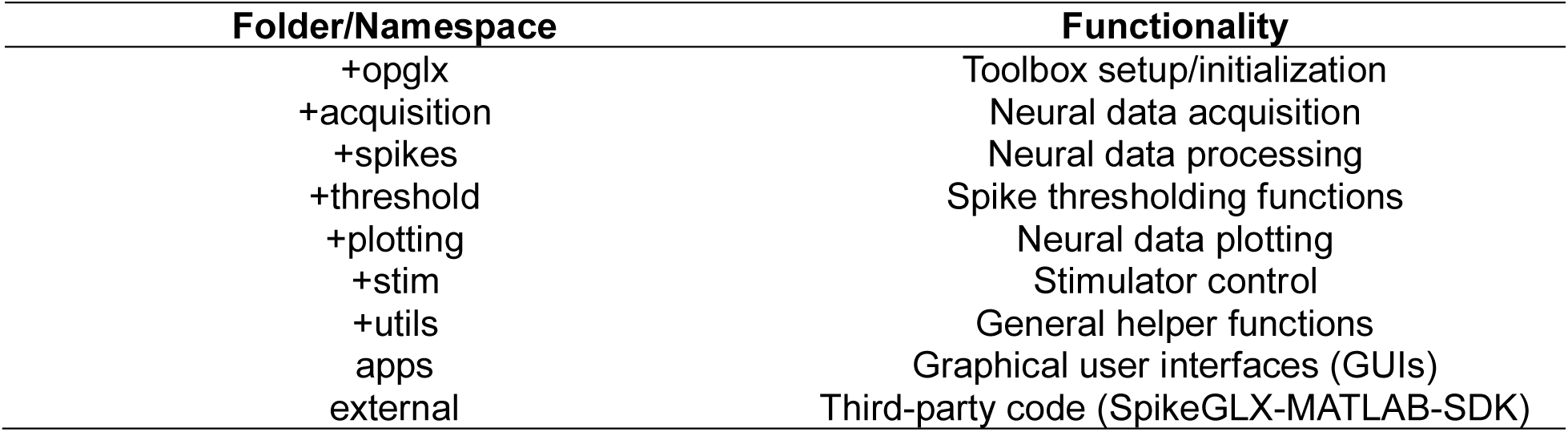
Toolbox code organization.

| Folder/Namespace | Functionality |
| --- | --- |
| +opglx | Toolbox setup/initialization |
| +acquisition | Neural data acquisition |
| +spikes | Neural data processing |
| +threshold | Spike thresholding functions |
| +plotting | Neural data plotting |
| +stim | Stimulator control |
| +utils | General helper functions |
| apps | Graphical user interfaces (GUIs) |
| external | Third-party code (SpikeGLX-MATLAB-SDK) |

#### Data Fetching

SpikeGLX maintains first in, first out (FIFO) buffers to store the most recent samples from acquisition streams. For a given NP probe, two AP-band stream buffers are available for sampling: 1) an 8-second raw data buffer, and 2) a configurable bandpass filtered, demultiplexed, global common average referenced (CAR) buffer of up to two seconds in length that runs in parallel with the raw stream. The *SpikeGL* API exposes functions to query the current sample count of a chosen stream (*GetStreamSampleCount*) and to retrieve data from a specified sample index (*Fetch*) or from the most recently available samples (*FetchLatest*).

Online processing of continuous data streams imposes the two primary constraints for a system utilizing the *SpikeGL* API:

1. contiguous samples must be retrieved, and
2. sample loss must be avoided or minimized.

For these reasons, OP-GLX exclusively utilizes *Fetch* rather than *FetchLatest*. Because *FetchLatest* does not allow for a current stream sample count to be directly specified, its use can result in overlapping samples (if time between consecutive calls is too short or the requested number of samples is too large) or missed samples (if time between consecutive calls is too long). In contrast, *Fetch* enables explicit sample count querying, provided that additional constraints are met:

1. *Fetch* calls must reference samples still present in the SpikeGLX buffer, and
2. *Fetch* calls must be adequately spaced to avoid saturation of the SpikeGLX command server.

For a given sample count *s0* returned by *GetStreamSampleCount*, a *Fetch* call requesting samples at *s0* must occur within the length of time that *s0* is present in the stream buffer (e.g. within 8 seconds for raw streams or up to two seconds for the filtered streams). Calling *Fetch* with an *s0* that is no longer present in the buffer results in a “FETCH: Too late” error.

Conversely, extremely high-frequency *Fetch* calls can result in freezing of the SpikeGLX console and stream viewer windows potentially leading to the acquisition crashing. Attempting to reestablish a connection will result in a “Receive timed out” error.

To operate within these constraints, OP-GLX implements the *SpikeFetcher* class, which serves as the core data fetching and scheduling component of the toolbox. A *SpikeFetcher* instance can be run in two modes: 1) *Continuous*, in which data are fetched and processed indefinitely until user termination, and 2) *Event*, in which data are fetched and processed around externally defined experimental events. In this section, the *Continuous* mode is described as this mode encompasses the core fetching and processing mechanisms used by OP-GLX. *Event* mode builds directly on this infrastructure and is detailed separately below. A sequence diagram illustrating the *Continuous* mode workflow along with GUI integration (detailed in a later section) can be seen in *Figure 2*.

**Figure 2:**
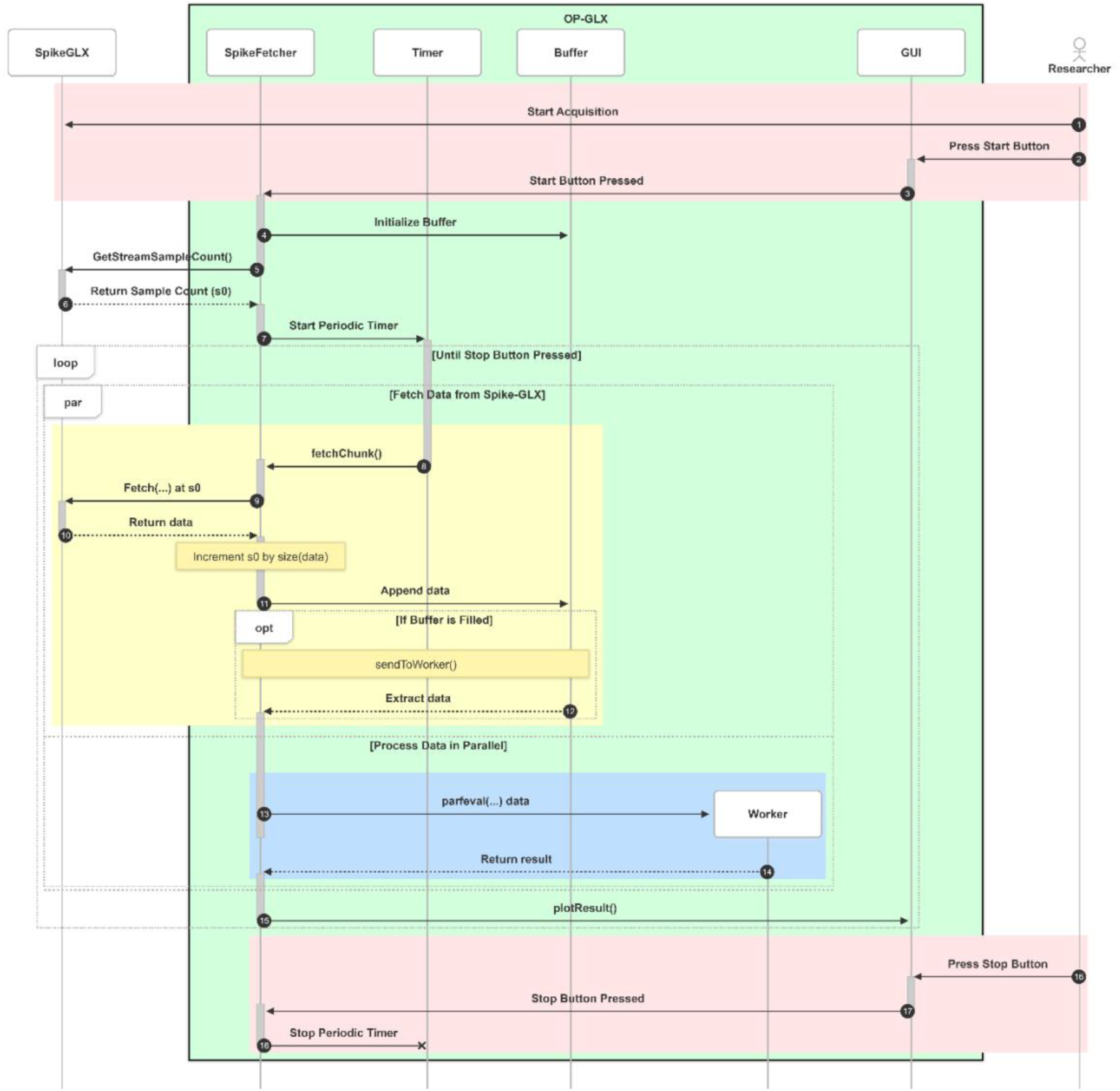
Sequence diagram of OP-GLX core functionality. Through GUI interaction, a researcher can start and stop data fetching from SpikeGLX via the OP-GLX *SpikeFetcher* class. Upon start, a MATLAB-side first-in, first-out (FIFO) buffer is initialized and the current SpikeGLX acquisition stream sample count is queried by API call *GetStreamSampleCount(…)* to set the initial fetch index (*s0*). A MATLAB timer is then started to periodically call *fetchChunk(…)* which requests available samples from SpikeGLX stream buffers starting from *s0* via API call *Fetch(…)*, updates *s0* based on the number of returned samples, and writes the data to the FIFO buffer. Once the buffer accumulates a user-specified window of data, *sendToWorker(…)* is called where the data is extracted and passed to a parallel worker for processing to allow data fetching to continue uninterrupted. When processing completes, results are returned to the main MATLAB session and the GUI plots are updated. This loop continues until the researcher presses the stop button which stops the periodic timer, halting data fetching and online processing.

To initiate *Continuous* mode operation, the *SpikeFetcher* method *start(fetchType)* is called with *fetchType* set to “Continuous”. Upon initiation, *SpikeFetcher* queries the current stream sample count with the API call *GetStreamSampleCount* and stores the returned value as the initial fetch index, *s0*. Rather than continuously polling SpikeGLX in a loop, *SpikeFetcher* uses a MATLAB timer to schedule periodic calls to its method *fetchChunk()* at a user-specified fetch interval. This interval is expressed as a fraction of the processing window length which is defined as the amount of data processed as a single unit. Each call to *fetchChunk()* does the following:

1. requests available samples from the SpikeGLX buffer starting at *s0* via *Fetch* API call,
2. updates *s0* based on the number of returned samples,
3. writes the returned data to a MATLAB-side FIFO buffer, and
4. checks if the buffer has filled to the user-specified processing window length

Through explicit control of fetch frequency and decoupling of fetch scheduling from processing, this approach prevents excessive API calls that could degrade SpikeGLX performance while ensuring retrieval of new samples occurs in a timely manner.

#### Windowing and Parallel Processing

To minimize blocking of *Fetch* API calls, which could lead to an *s0* falling outside of the SpikeGLX stream buffer, data processing is performed using parallel workers. Specifically, OP-GLX utilizes MATLAB’s thread-based environment via *parpool(‘Threads’)* which allows workers to share memory with the main MATLAB session, avoiding the potential overhead associated with copying large data arrays to other processes.

When the number of samples accumulated in the MATLAB-side buffer reaches the user-specified processing window length, the *SpikeFetcher* method *sendToWorker()* is called at the end of the current *fetchChunk()* execution. Because of the limited precision and variability of MATLAB timers, there is no guarantee that the next *Fetch* API call will return the same number of samples as the previous call. Explicitly setting a window length ensures that data is processed with a static size allowing other variables, such as time arrays and plotting objects, to be pre-allocated. Once filled, data is extracted from the buffer based on the window length and a processing function is assigned based on the type of analysis or plot the user has selected.

The data and function are then passed to a parallel worker for processing using the MATLAB *parfeval* function. Finally, a plotting function is assigned to update plots in the main MATLAB session once the worker finishes processing.

Presently, OP-GLX provides four processing functions (found in the +spikes namespace) that support GUI visualization (detailed in a later section):

1. *rasterSpikes*, which returns rasterized spike times from all channels
2. *getSpikeWaveforms*, which returns AP-band traces and aligned spike waveforms
3. *getSpikeFiringRates*, which computes time-binned firing rates across channels
4. *pcaSpikes*, which computes PCA on time-binned firing rates and returns the first three principal components over time.

All four functions rely on an underlying spike detection function which identifies threshold crossing events across all channels of a NP probe. OP-GLX includes two functions: *detectSpikes* and *detectSpikesLW*, a lightweight version of the former. Spike detection is implemented with the following general procedure:

1. Noise standard deviation (SD) is estimated per channel via user-selectable estimation method.
2. Candidate spikes are identified via threshold crossing using a user-specified multiple of the noise estimate.
3. Removal of artifacts not eliminated via CAR by blanking candidate “spikes” that occur on more than a user-specified fraction of channels simultaneously (a simplified approach to the *-gfix* option from the offline CatGT tool[14]).
4. Candidates are confirmed as spikes if threshold is crossed for a specified number of consecutive samples (stay-below criteria).

At present, the supported noise SD estimation methods are median absolute deviation (MAD) [15], [16] which is located in the *+threshold* namespace and MATLAB’s built-in SD function *std*. MAD implementation *madEstimation* uses MATLAB’s *mad* function multiplied by the standard normal scaling factor (1.4826). When fetching from the filtered stream buffers, CAR-applied data is returned zero-median. In this case, *madEstimationZeroMedian* can be used to avoid redundant median calculations. Additionally, custom estimation functions can be written following the format of the existing functions and placed in the *+threshold* namespace.

Importantly, the implementation of step 4 returns spike times corresponding to the first sample which meets the stay-below criteria and does not guarantee a local minimum. In *detectSpikes*, a final step is performed to search for and adjust spike times to the local minimum of each threshold crossing. The search is conducted using a look-ahead window which can be adjusted by the user. Additionally, small above-threshold gaps may be present between below-threshold samples resulting in either two “spikes” being detected or a spike being missed. To handle this, *detectSpikes* fills in one-sample gaps by default or set by the user. In cases where it is not critical that spike times reflect the minimum of an action potential, *detectSpikesLW* can be used which returns spike times as the first sample meeting the stay-below criteria.

Detected spike times and corresponding channel indices are returned for further processing. The detection algorithm is fully vectorized and, along with the four processing functions and noise estimation methods, fully supported in MATLAB’s thread-based environment.

#### Stimulus Control and Event Fetching

During recordings involving experimental events, such as somatic [17], visual [18], [19], or electrical stimuli [20], [21], [22], it can be helpful to analyze and visualize brain responses while the experiment is ongoing. The *Event* mode of *SpikerFetcher* provides this functionality when used with the toolbox’s *StimulationInterface* class (*toolbox/+stim*). StimulationInterface is an abstract class with properties, methods, and events that can be used by a *SpikeFetcher* instance or within the GUI. A custom class is written which inherits and defines the properties and methods to be used with a specific stimulator.

An example stimulator class is provided, the *ArduinoStimulator* interfaces an Arduino microcontroller to control a vibration motor for delivering vibrotactile stimuli. In this example setup, a pin on the Arduino is connected to a digital input of a National Instruments (NI) board configured for SpikeGLX acquisition. The duration and amplitude of the vibrotactile stimulus can be set by *updateParameters()* and delivered via *deliverStimulus()*. When delivered, the Arduino pin goes high for the duration of the stimulus and is acquired on one bit of the digital input channel.

Once the stimulator is setup, a MATLAB event listener is attached to call *deliverStimulus()* when a *SpikeFetcher* notifies “DeliverStimulus”. *SpikeFetcher* can now be run in *Event* mode, which starts a MATLAB timer to call the *findEvent()* method and notifies the stimulator to deliver the stimulus. In *findEvent()*, the NI acquisition stream is queried with the *SpikeGL* API to scan for, in this example, the rising edge of the stimulus event, and is called periodically by a user-specified frequency until an event is found. Once the event is found, the *fetchEvent()* method is called which:

1. Stops and reassigns the timer to call *fetchChunk()* as in *Continuous* mode.
2. Maps the identified event sample count of the NI stream to the NP data stream using the API call *MapSample*
3. Set the running *s0* counter to the mapped sample minus a user-specified pre-stimulus window
4. Starts the timer which calls *fecthChunk()* until the MATLAB-side buffer is filled to the user-specified pre-stimulus, stimulus, and post-stimulus windows length

When all samples around the event are fetched from SpikeGLX, the MATLAB timer is stopped, and the data is processed and displayed in a figure window or GUI with vertical lines marking the start and stop of the event.

#### GUI Design, User Interaction, and Visualization

While *SpikeFetcher* can be run independently in the MATLAB command window or via scripts, OP-GLX is packaged with a GUI that provides integrated control of data fetching, processing, and visualization (Fig. 3a). After beginning a SpikeGLX acquisition (Fig. 3b), a user can open the GUI (*toolbox/apps/opglx.mlapp*) and press the *Initialization* button which queries parameters of the run. Once initialized, a user can select to start a *Continuous* or *Event* (if a stimulator is enabled) run. Most parameters can be changed while in *Continuous* mode and are applied immediately via the *ParamaterManager* handle class. Switching the plot tabs at the top of the app similarly switches the type of analysis to be performed and takes effect at the next *sendToWorker()* call.

**Figure 3:**
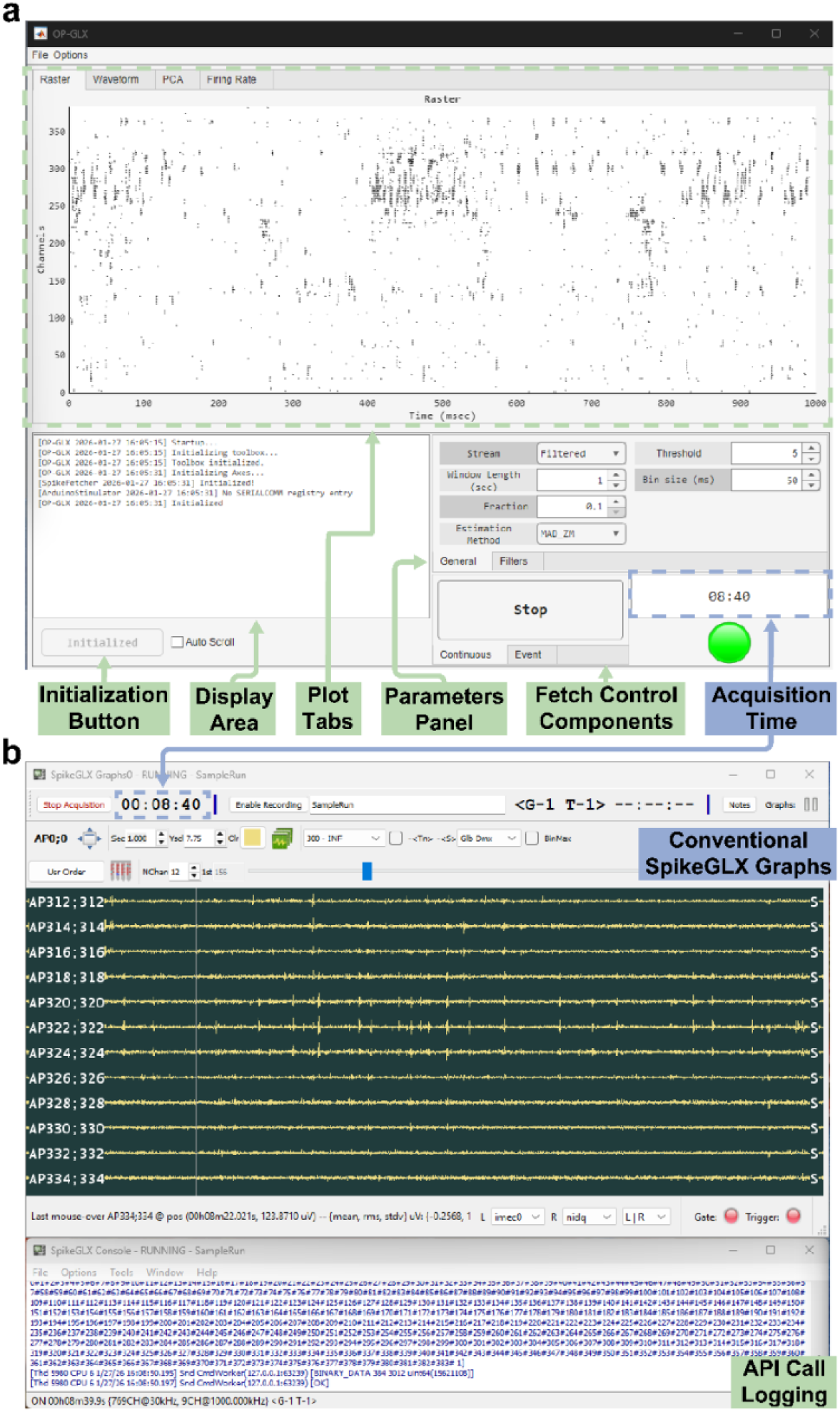
OP-GLX graphical user interface (GUI) operating during SpikeGLX acquisition. **a**) The OP-GLX toolbox is packaged with a MATLAB-based GUI that utilizes *SpikeFetcher* for online processing and visualization of SpikeGLX NP data streams. The GUI provides controls for initialization, data fetching, setting parameters, and selection of plot tabs to switch between raster display, waveform display, PCA trajectory, and firing-rate estimates. When set to *Continuous* mode, data fetching, processing, and plotting will continue until the stop button is pressed. **b**) SpikeGLX graph viewer (top) and console (bottom) running on the same system showing conventional AP-band displays and command server/API logging. SpikeGLX remains responsible for data acquisition while OP-GLX interfaces with SpikeGLX via the MATLAB API to enable online, probe-wide analysis and visualization during experiments.

### Implementation and Performance

A critical feature of OP-GLX is to reliably schedule fetching, processing, and plotting of data while remaining within and at the front of the SpikeGLX buffers. To measure and quantify the performance and stability of the toolbox in a real-time setting, a sample NP recording was simulated in SpikeGLX and a *SpikeFetcher* instance was set to fetch from the filtered stream in *Continuous* mode for multiple fetch lengths and processing window lengths. For each fetch length and processing window length combination, *SpikeFetcher* collected performance metrics for 60 seconds. Fetch lengths equal to or below a given window length were tested. In all cases, online processing was set to compute and plot rasterized spike times using *rasterSpikes* which calls *detectSpikes* (Fig. 4) or *detectSpikesLW* (Supp. Fig. 1).

**Figure 4:**
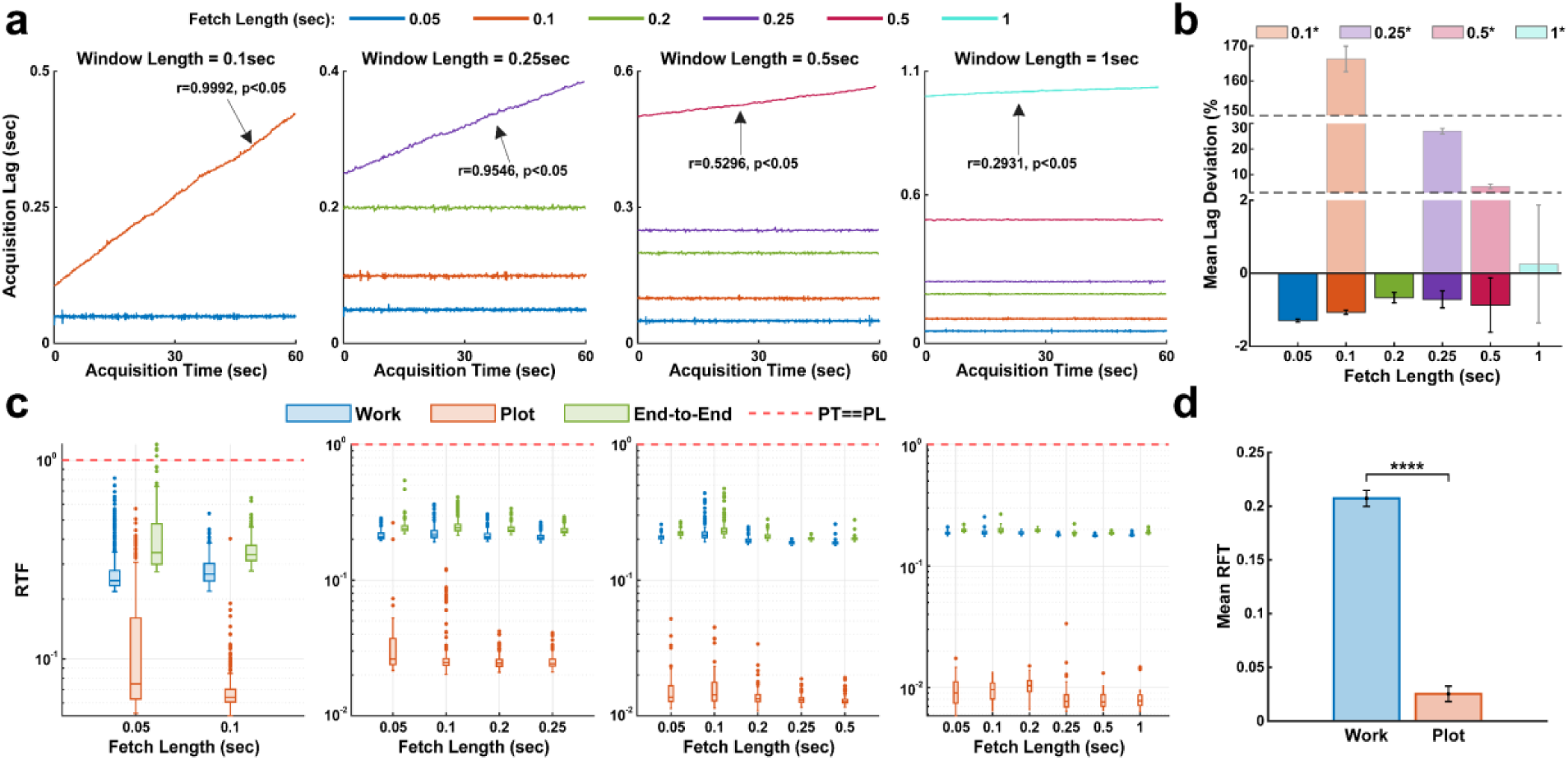
Performance metrics of OP-GLX *detectSpikes* and raster plotting during *Continuous* mode operation. **a**) Acquisition lag as a function of acquisition time across 60-second runs for multiple fetch lengths and processing window lengths. Black text arrows indicate a significant positive correlation between acquisition time and acquisition lag (Pearson’s correlation coefficient). **b**) Acquisition lag deviation from nominal fetch length (y==0) for each fetch length averaged across window lengths. Dark bars indicate performance runs where fetch length is less than the window length. Light bars indicate performance runs with positive correlation from (**a**) where fetch length is equal to window length as represented by * in the legend. Error bars indicate SEM. **c**) Real-time factor (RTF) computed for processing (Work), plotting (Plot), and total (End-to-End) times. Dashed red line indicates RTF of 1, when the processing time (PT) equals the processing length (PL). Boxes extend from 25th to 75th percentiles with horizontal lines indicating median. Filled circles indicate outliers identified as values 1.5x(interquartile range) away from 25th and 75th percentiles. Whiskers correspond to the maximum and minimum non-outlier values. Columns indicate window lengths as in (**a**). **d**) Mean Work and Plot RTF averaged across all fetch length and window lengths combinations (n=17 runs). Error bars indicate SEM. ****: p<0.0001 (two-tailed paired t-test).

#### Neural Dataset

The data used for assessing the toolbox consisted of NP recordings from follow-up experiments of a previous study [23]. In that study, the data were collected during an acute implantation of an NP probe (Neuropixels 1.0, IMEC, Belgium) in a urethane-anesthetized (1.2 g/kg, IP) rat (Long-Evans, male, 2–3-month-old). During the experiment, the rat was stabilized in a stereotaxic frame with ear bars, and probe trajectory and craniotomy location were planned using the Pinpoint trajectory planning software [24] with the Waxholm rat atlas to target the somatosensory cortex (S1) and ventral posterolateral thalamic nucleus (VPL). The probe insertion coordinates were (AP: -0.92 mm; ML: -2.68 mm; azimuth: -169°; elevation: 65°) relative to bregma. After dura removal and securing a ground screw near the craniotomy, a stereotaxic micromanipulator (M3-LS-3.4-15-XYZ-MPM-Inverted, New Scale Technologies) was used to insert the NP probe 7.81 mm into the brain before initiating neural recordings.

#### Acquisition Lag

To quantify the delay between a *fetchChunk()* call and the current acquisition time, acquisition lag is measured. Acquisition lag is defined as the difference between the current acquisition time and the current *s0* to request samples from with the *Fetch* API call. At the top of *fetchChunk()*, *GetStreamSampleCount* is queried directly before the *Fetch* API call to provide a value of the current acquisition time. As the MATLAB timer schedules *fetchChunk()* calls periodically based on the fetch length, it would be expected that the acquisition lag should be approximately equal to the set fetch length. Additionally, the acquisition lag should remain constant (zero slope) over the course of the acquisition. An increase in acquisition lag over time indicates that *SpikeFetcher* is falling behind the SpikeGLX stream buffers which will eventually lead to *s0* requests that are no longer in the buffers.

Over the course of 60 second runs, acquisition lag remained stable when fetch length was not equal to window length (Fig. 4a, Supp. Fig. 1a). In these cases, across different window lengths, acquisition lag was less than the nominal fetch length by 1.29 ± 0.04%, 1.06 ± 0.06%, 0.67 ± 0.14%, 0.71 ± 0.74%, and 0.87 ± 0.74% for fetch lengths of 0.05, 0.1, 0.2, 0.25, and 0.5 sec, respectively (Fig. 4b, dark bars). Similar acquisition lag trends were found when *detectSpikesLW* was used (Supp. Fig. 1b). This indicates that fetching remained stable over time, remaining well within the SpikeGLX stream buffers. When fetch length was equal to the window length, a steady increase in acquisition lag was observed, with significant positive correlation for both *detectSpikes* and *detectSpikesLW* (Fig. 4a, Supp. Fig. 1a, text arrows). When running *detectSpikes*, for fetch lengths of 0.1, 0.25, 0.5, and 1 sec, acquisition lag was observed to be greater than the nominal fetch length by 166.29 ± 3.67%, 27.00 ± 1.11%, 5.44 ± 0.8%, and 0.26 ± 1.61%, respectively (Fig. 4b, light bars). For *detectSpikesLW*, three of the four cases (0.1, 0.25, and 0.5 sec) found acquisition lag to be greater than the nominal fetch length with the fourth case (1 sec) marginally below the nominal value (Supp. Fig. 1b, light blue bar).

While fetch errors did not occur during the 60-second acquisition tests in these four cases, it is expected that lag will continue to accumulate until reaching the stream buffer length and will eventually fail with a “FETCH: Too late” error.

#### Real-Time Factor (RTF)

We calculated real-time factor (RTF) to quantify the processing performance of the parallel workers. RTF was computed as the ratio of processing time (PT), defined as the elapsed wall-clock time required for a parallel worker to complete computation, to processing length (PL), defined as the duration of neural data being processed. An RTF of 1 indicates that processing time keeps pace with the incoming data, while values below 1 indicate real-time capability with available computational margin. Timestamps were recorded immediately before data were dispatched to a parallel worker, upon worker completion, and after plot updates in the main MATLAB session.

Using these timestamps, RTF was computed separately for the time spent performing data processing (“Work”), updating the visualization object (“Plot”) and total end-to-end latency from dispatch to plot completion (“End-to-End”). For all tested combinations of fetch length and processing window length, RTF values for all three PT stages remained below 1 (Fig. 4c, Supp. Fig. 1c). This indicated that probe-wide spike detection and rasterization consistently completed faster than the duration of the processed data window, confirming that OP-GLX was able to sustain real-time operation under the tested conditions.

Across window and fetch lengths, Work RTF was significantly greater than Plot RTF (mean ± SD: 0.1820 ± 0.0142; 95% CI [0.1747, 0.1893], p < 0.0001, Fig. 4d), indicative of the computational cost of spike detection and rasterization relative to the graphical updates. Similar RTF trends were observed when *detectSpikesLW* was utilized (Supp. Fig. 1d). Plotting, while faster on average, exhibited greater variance across fetch lengths and trials, likely reflecting variability in MATLAB’s graphics rendering and event queue scheduling. Across conditions, End-to-End RTF remained well below 1 (mean ± SD: 0.2326 ± 0.0584; 95% CI [0.2025, 0.2626]; p < 0.0001, two-tailed one-sample t-test against µ = 1) indicating that combined processing and visualization latency does not accumulate over time. Notably, as the PT measurements were not taken in reference to the current acquisition time, the lag accumulation observed in Fig. 4a was not present in RTF.

Together, these results demonstrate that OP-GLX provides sufficient computational performance for real-time analysis and visualization of NP data across a range of processing window sizes and fetch lengths, while maintaining stable end-to-end latency necessary for online experimental monitoring.

#### Comparison to Offline Analysis

Finally, to assess the ability of OP-GLX to capture meaningful information about the neural acquisition, online processing methods were compared to offline spike-sorting using Kilosort [25]. Using a 30-second recording, *detectSpikes* was run using a threshold of 4x (and default values for all other parameters) on 0.5-second chunks of data to return channel-based spikes times across the entire recording. Spike-sorting was performed on the same recording and, to provide a direct comparison to the channel-based method, spike times from identified single-units were labelled based on the peak channel of the given single-unit (Fig. 5a). Across the recording, *detectSpikes* detected 157,807 spikes originating from 376 channels and Kilosort extracted 48,680 from 156 single-units mapped to 152 channels.

**Figure 5:**
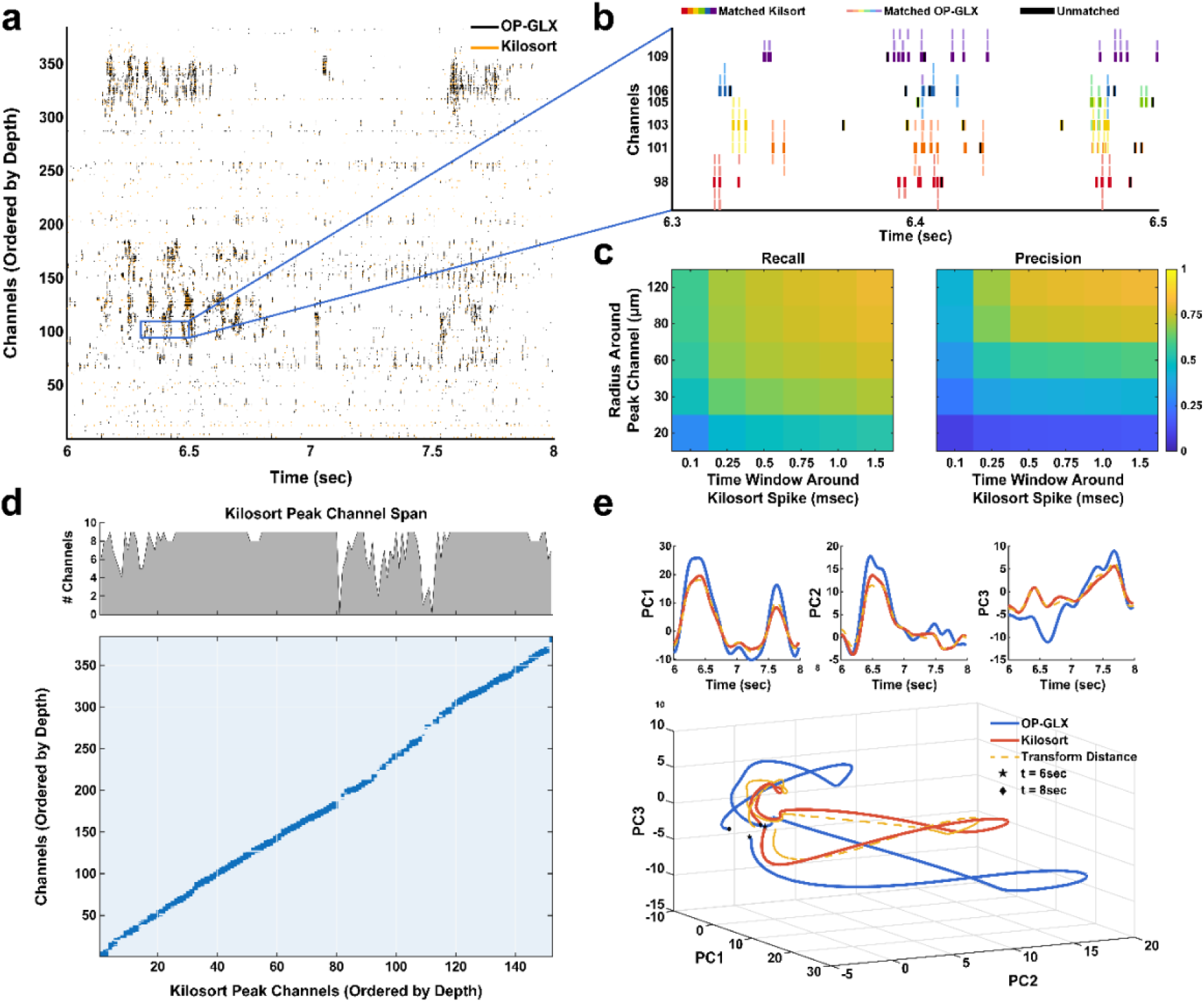
Evaluation of OP-GLX processing methods. **a**) Raster plot of spike times detected via channel-based *detectSpikes* function (black) and from offline spike-sorting with Kilosort (orange). Kilosort spikes from single-units were labelled corresponding to the peak channel of each spike’s unit. **b**) Visualization of the spike-matching method used to match OP-GLX spikes to Kilosort spikes for channel radius of 80 µm and time window of 1 msec. Thick, colored ticks indicate Kilosort spikes that were matched by at least one *detectSpikes* spike. Thin, colored ticks represent *detectSpikes* spikes that were matched to a Kilosort spike. Black ticks indicate Kilosort spikes that were not matched with a *detectSpikes* spike. **c**) Heatmaps of recall (fraction of Kilosort spikes detected via *detectSpikes*) and precision (fraction of OP-GLX spikes matched to Kilosort spikes) for multiple time window lengths and channel radii. **d**) Biadjacency matrix of spike matching for channel radius of 80µm and time window of 1ms. Blue squares indicate that at least one *detectSpikes* spike from a given channel (y-axis) matched with a Kilosort spike at its unit’s peak channel (x-axis). Bar plot represents the number of channels with at least one spike that matched a given Kilosort peak channel. For a radius of 80µm, a maximum of 9 channels around a Kilosort channel could contain matches based on probe mapping and geometry. **e**) PCA trajectories during the time-course in (**a**) of firing rates estimated from *detectSpikes* or Kilosort spikes. The first three principal components of *detectSpikes*-derived trajectories were transformed into Kilosort-derived trajectories following Procrustes analysis.

Two key metrics were calculated: 1) The fraction of Kilosort spikes that were detected by *detectSpikes* (recall), and 2) the fraction of *detectSpikes* spikes that matched with a Kilosort spike (precision). Matches were considered when a *detectSpikes* spike from a channel within a specified radius around each Kilosort peak channel fell within a specified time window (match window) around a Kilosort spike (Fig. 5b). Recall and precision, measured across multiple radii (20, 30, 60, 80, 120 µm) and match window lengths (0.1, 0.25, 0.5, 0.75, 1.0, and 1.5 msec) were found to increase with increasing radius and match window length (Fig. 5c). Highest values were found to be 0.8039 and 0.8091 for recall and precision, respectively at a radius of 120 µm and match window of 1.5 ms. However, as the radius and match window are relatively large, a radius of 80 µm and match window of 1 ms were used in the remaining analyses (recall - 0.7707; precision – 0.7582). Matching was further characterized by measuring the number of channels with at least one spike that matched to a Kilosort spike per Kilosort peak channel (Fig. 5d). Averaged across all combinations of radii and match windows, a given Kilosort peak channel matched with *detectSpikes* spikes from 5 ± 3 channels. While spike-sorting may not be considered a true “ground-truth”, these results indicate that *detectSpikes* reliably detects “ground-truth” spikes based on offline analysis.

Lastly, population activity representations using principal component analysis (PCA) were examined. Specifically, PCA was performed on z-scored firing rates estimated from time-binned spikes and smoothed using a gaussian kernel. Trajectory similarity of the first three principal components derived from *detectSpikes* spikes or Kilosort spikes was next assessed using Procrustes analysis without reflection (Fig. 5e, Supp. Fig. 2a). Procrustes squared distance between PCA trajectories across firing rate estimation parameters was found to be low, within a range of 0.08-0.15 (Supp. Fig. 2b) indicating lower dimensional representations of neural activity is preserved by OP-GLX processing methods.

## Discussion and Conclusion

The widespread adoption of high-density NP probes has generated a growing demand for software systems that can operate on high-bandwidth neural data streams in real time. While SpikeGLX provides a robust, low-latency platform for data acquisition, its online processing and plotting capabilities are intentionally limited to prioritize acquisition stability. OP-GLX was developed to address this gap by providing a flexible, MATLAB-based toolbox for enabling real-time fetching, processing, and visualization of NP data that seamlessly integrates with existing SpikeGLX workflows.

A central design challenge addressed by OP-GLX was to maintain reliable, online performance within the constraints imposed by the SpikeGLX ecosystem. These constraints required that API data requests must be carefully scheduled to avoid both 1) falling behind the SpikeGLX stream buffers when calls are too infrequent (“FETCH: Too late”) and 2) saturating the command server when calls are too rapid. OP-GLX addressed these constraints by decoupling data request scheduling from data processing using MATLAB timers and buffers, fixed-sized processing windows for deterministic downstream computation, and parallel processing to prevent blocking of data requests. Performance testing confirmed that when fetch lengths were chosen to be shorter than the processing window length, acquisition lag remained stable and bounded over time. This confirms that OP-GLX can remain reliably synchronized with the SpikeGLX stream buffers during sustained operation.

A critical architectural decision that enabled real-time performance was the use of parallel processing with MATLAB’s thread-based workers. By offloading spike detection and signal processing to shared-memory worker threads, OP-GLX minimized blocking of *Fetch* API calls and avoided the overhead involved with inter-process data copying. RTF measurements indicated that probe-wide spike detection and visualization consistently completed faster than the duration of the processing window, demonstrating that OP-GLX can sustain real-time operation when analyzing all channels of an NP probe simultaneously. This performance margin can allow users to trade computational headroom for more sophisticated analyses or shorter processing windows when lower latency is desired.

Despite these demonstrated capabilities, OP-GLX has important limitations. Primarily, OP-GLX performs *indirect* streaming of neural data, relying on the SpikeGLX MATLAB API rather than directly accessing the acquisition buffers. Due to this indirect streaming, the minimum achievable end-to-end latency of OP-GLX is ∼6.5ms [13], which is defined by the measured round-trip latency of the SpikeGLX API. The < 10 milliseconds latency is acceptable for most purposes such as online monitoring of neural data during experimental events or behavioral tasks [17], [18], [19], [22], [26], [27], [28], and closed-loop experiments which necessitate data binning on the order of tens of milliseconds [20], [21], [29], [30]. However, OP-GLX may not be ideal for applications requiring precise millisecond to sub-millisecond feedback such as circuit-level investigations of synaptic plasticity [31], [32], [33].

Additional limitations arise from the design choice of prioritizing performance and usability over computationally intensive processing methods. The implemented spike detection algorithm in OP-GLX was intentionally lightweight and performed at the channel-level. Although sufficient for online firing-rate estimation and population-level analyses, this approach did not employ spike-sorting methods for resolving single- and/or multi-units in real-time. Furthermore, the current implementation was designed around single-probe operation. While adding probe selection functionality may be relatively straightforward, simultaneously fetching from multiple NP probes will require additional coordination and optimization of fetch scheduling, parallel worker allocation, and memory management to maintain real-time performance.

Several future directions could substantially extend the utility of the OP-GLX toolbox. Incorporating online spike sorting would enable single- and multi-unit identification during experiments. Extending our architecture to support multiple probes would broaden its use in large-scale recording paradigms that are increasingly common in systems neuroscience [4], [34], [35]. Finally, translating or reimplementing core components in Python or C/C++, either by the SpikeGLX-CPP-SDK [36] or a lightweight server-client architecture, could reduce latency, remove MATLAB-specific overhead and dependencies, and enable tighter integration with the SpikeGLX control systems.

At present, OP-GLX provides a practical and customizable toolbox for online processing and visualization of NP data acquired by SpikeGLX. By explicitly addressing the timing and buffering constraints of SpikeGLX and its MATLAB API, and by enabling parallel computation in MATLAB, OP-GLX provides stable, online analyses across all channels of a Neuropixel probe during ongoing experiments. As high-density neural recording technologies continue to be adopted, tools such as OP-GLX will play an important role in bridging the gap between data acquisition and analysis, facilitating more informed experimental control, and furthering progress in systems neuroscience and neural engineering.

---

**Supplementary Figure 1:**
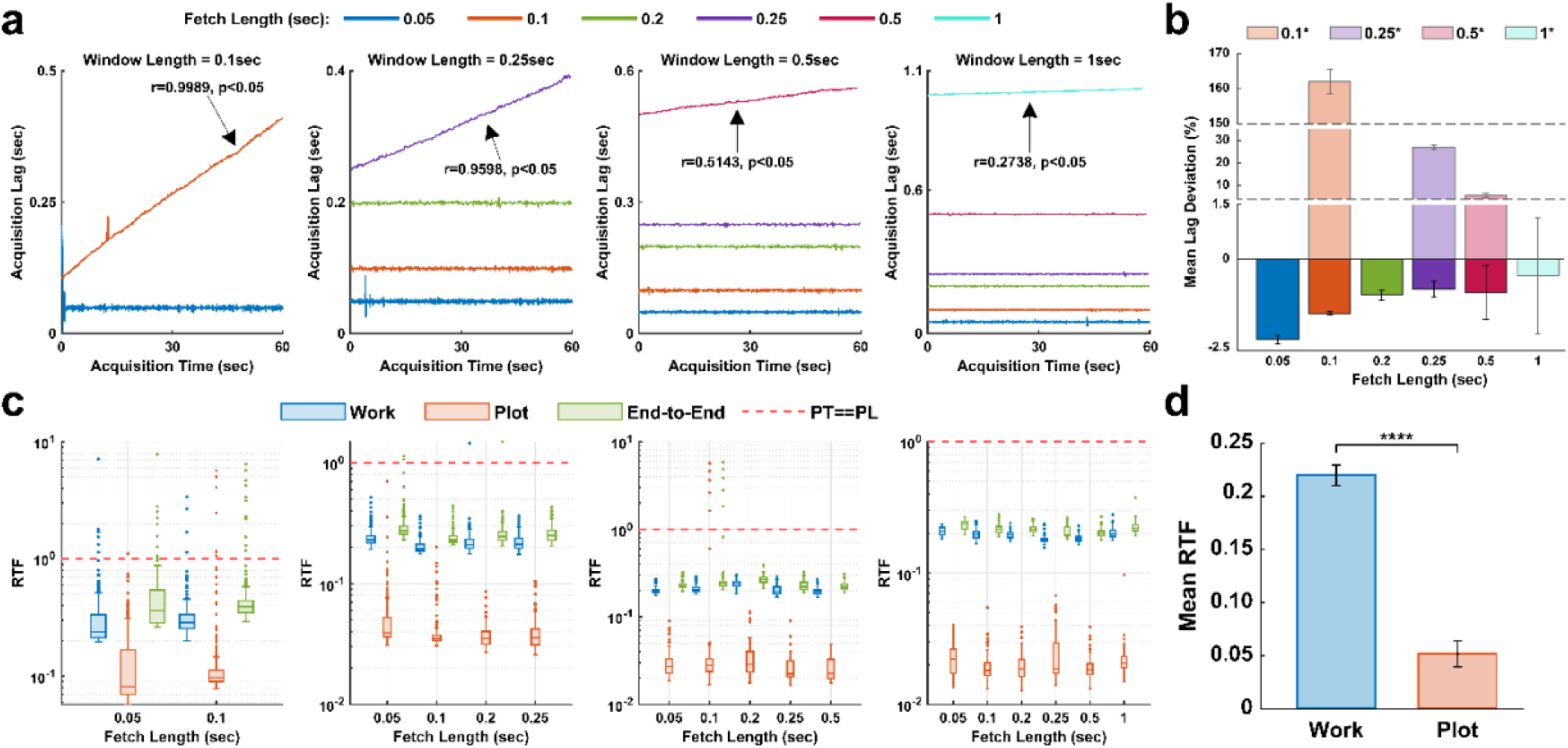
Performance metrics for *detectSpikesLW*. **a**) Same analysis of acquisition lag as in Fig. 4a but for light-weight spike detection function *detectSpikesLW*. **b**) Same analysis acquisition lag deviation as in Fig. 4b for *detectSpikesLW*. **c**) Same analysis of Work, Plot, and End-to-End RTF as in Fig. 4c for *detectSpikesLW*. **d**) Same analysis of condition-averaged Work and Plot RTF as in Fig. 4d for *detectSpikesLW*.

**Supplementary Figure 2:**
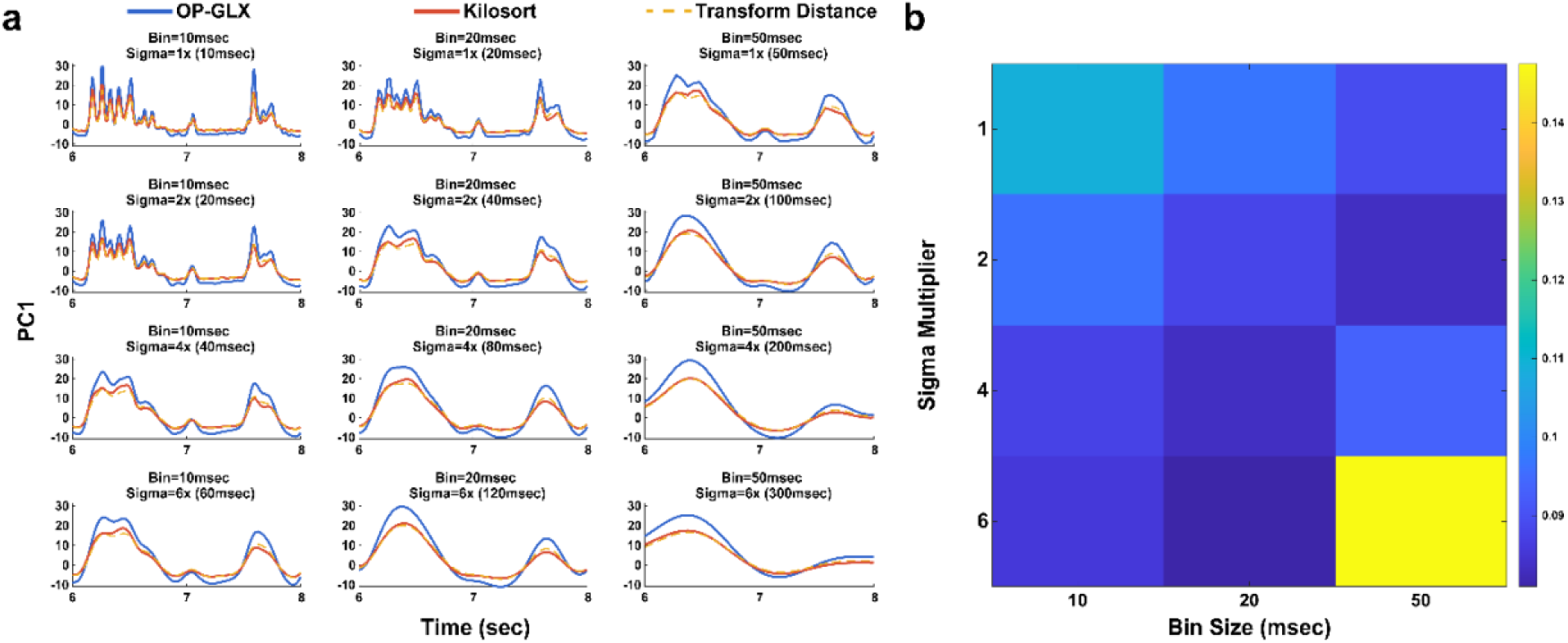
Procrustes distance of PCA trajectories across firing rate estimation parameters. **a**) Sample trajectories of PC1 and resulting Procrustes transformation across the time range as in Fig. 5a and 5e for firing rates estimating for multiple bin widths (10, 20, 50 msec, columns) and sigma values for the Gaussian kernel (1, 2, 4, 6 x bin width, rows). **b**) Heatmap of Procrustes squared distance for firing rate parameter combinations as in (**a**) computed across the entire recording length.

<u>Supplementary Video 1: Example usage of OP-GLX alongside SpikeGLX.</u> After configuring parameters and beginning an acquisition in SpikeGLX, the OP-GLX app can be run. Following initialization, OP-GLX fetches incoming data from SpikeGLX and performs multiple processing and plotting operations in real-time.

